# Decreased sarcomeric mitochondrial creatine kinase 2 impairs skeletal muscle mitochondrial function independently of insulin action in type 2 diabetes

**DOI:** 10.1101/2024.01.11.575194

**Authors:** David Rizo-Roca, Dimitrius SPSF. Guimarães, Logan A. Pendergrast, Nicolas Di Leo, Alexander V. Chibalin, Salwan Maqdasy, Mikael Rydén, Erik Näslund, Juleen R. Zierath, Anna Krook

## Abstract

Plasma creatine levels are associated with risk of type 2 diabetes. Since skeletal muscle is the main disposal site of both creatine and glucose, we investigated the role of intramuscular creatine metabolism in the pathophysiology of insulin resistance in type 2 diabetes. We report in men with type 2 diabetes, plasma creatine levels are increased, while intramuscular phosphocreatine content is reduced. These alterations are coupled to reduced expression of sarcomeric mitochondrial creatine kinase 2 (*CKMT2*). In C2C12 myotubes, *Ckmt2* silencing reduced mitochondrial respiration, membrane potential, and glucose oxidation. Electroporation-mediated overexpression of *Ckmt2* in skeletal muscle of high-fat diet-fed male mice increased mitochondrial respiration, independent of creatine availability. Thus, beyond the canonical role of CKMT2 on creatine phosphorylation, we reveal a previously underappreciated role of CKMT2 on mitochondrial homeostasis, independent of insulin action. Collectively, our data provides functional evidence into how CKMT2 mediates mitochondrial dysfunction associated with type 2 diabetes.

## INTRODUCTION

Insulin resistance associated with type 2 diabetes is characterized by alterations in energy metabolism. Since intracellular storage and transport of ATP is limited, high-energy demanding tissues, such as brain and skeletal muscle utilise creatine kinases to convert creatine into phosphocreatine. Creatine can be both efficiently stored and transported across the cell membrane and is therefore an important energy shuttle. Thus, creatine metabolism can couple mitochondrial energy production with cytosolic energy demands^1^.

Alterations in enzymes controlling creatine metabolism have emerged as multifaceted players in various pathophysiological contexts. For example, creatine metabolism mediates macrophage immune response^2^ and contributes to the metabolic rewiring observed in cancer cells^3^. In white adipose tissue, impaired phosphocreatine metabolism contributes to the development of a proinflammatory environment, linking alteration in creatine metabolism to obesity and metabolic disease^4^. Notably, increased circulating levels of creatine in plasma is associated with a higher risk of type 2 diabetes incidence in men^5^. Thus, correlative evidence suggests alterations in creatine metabolism may be involved in the development of metabolic dysfunction.

Perturbations in creatine metabolism stem from changes in the expression and activity of the creatine transporter and members of the creatine kinase family, proteins responsible for the membrane transport and interconversion of creatine into phosphocreatine. Creatine kinases exhibit tissue-specific and discrete intracellular localisation^6^. In heart and skeletal muscle, the creatine kinase M-type isoenzyme is found in the cytosol, whereas the creatine kinase S-type (also known as sarcomeric mitochondrial creatine kinase 2, sMtCK or CKMT2) is located in the intermembrane mitochondrial space. CKMT2 exists as a homo-octamer, consisting of four dimers that serve as the stable structural units^7^. CKMT2 functionally co-localizes with the adenine nucleotide translocator (ANT) to rapidly hydrolyse ATP into ADP to phosphorylate creatine^8^. Thus, given this tissue-specific distribution and intracellular localization, CKMT2 is a key regulator of oxidative phosphorylation and mitochondrial respiration in skeletal muscle^9^. However, the precise role for this enzymatic machinery in skeletal muscle metabolism in type 2 diabetes is unexplored. Skeletal muscle, is a primary tissue for both glucose^10^ and creatine disposal^11^, and therefore emerges as a crucial organ for investigating the role of creatine metabolism in the development of insulin resistance in type 2 diabetes

Herein, we investigated whether creatine metabolism and CKMT2 content are altered in skeletal muscle from men with type 2 diabetes. We also determined how modulation of creatine metabolism impacts metabolic health and mitochondrial function. By combining clinical and experimental approaches, we provide evidence that perturbations in CKMT2 content contributes to the impaired skeletal muscle mitochondrial homeostasis characteristic of type 2 diabetes.

## RESULTS

### Skeletal muscle creatine metabolism is altered in men with type 2 diabetes and linked to aberrant glucose control

To determine whether creatine metabolism is perturbed in skeletal muscle in type 2 diabetes, we collected plasma samples and *vastus lateralis* muscle biopsies from men with normal glucose tolerance or type 2 diabetes (**Figure 1A**. Clinical characteristics of the study participants in **Table 1** and **2**). We found that circulating fasting levels of creatine were increased in plasma from men with type 2 diabetes (**Figure 1B**) and were negatively correlated with the expression of the creatine transporter, *SLC6A8* in skeletal muscle (**Figure 1C-D**). Although the intramuscular content of creatine was higher in men with type 2 diabetes (**Figure 1E**), phosphocreatine levels were reduced (**Figure 1F**), and these levels were associated with the expression of the mitochondrial creatine kinase 2, *CKMT2* in skeletal muscle (**Figure 1G-H**). The intramuscular phosphocreatine level was inversely correlated with fasting blood glucose, with opposing trends in men with type 2 diabetes as compared to men with normal glucose tolerance (**Figure 1I**). Similarly, lower skeletal muscle *CKMT2* mRNA levels were associated with higher post-OGTT circulating insulin in men with normal glucose tolerance (**Figure 1J**) and higher haemoglobin A1c in men with type 2 diabetes (**Figure 1J-K**). While *CKMT2* correlated positively with the hip diameter in men with normal glucose tolerance, it associated with larger waist-to-hip ratio in men with type 2 diabetes (**Figure 1J**). The levels of creatine precursors in plasma were largely unaltered (**Figure S1A-E**), except for glycine, which was decreased in men with type 2 diabetes (**Figure S1A**). As BMI was higher in men with type 2 diabetes, we also examined whether BMI *per* se explained the relationship between creatine metabolism and clinical features. Except for the positive correlation between BMI and levels of plasma creatine levels (**Figure S1F**), BMI did not associate with either creatine/phosphocreatine or gene expression in skeletal muscle (**Figure S1F-J**). With the exception of intracellular skeletal muscle creatine, comparisons between diagnosis groups remained significant after adjusting for BMI. Overall, our data provide evidence that key elements of creatine metabolism are disrupted in skeletal muscle of men with type 2 diabetes. Thus, alterations in creatine metabolism may be linked impaired glucose metabolism in type 2 diabetes.

**Figure 1.**
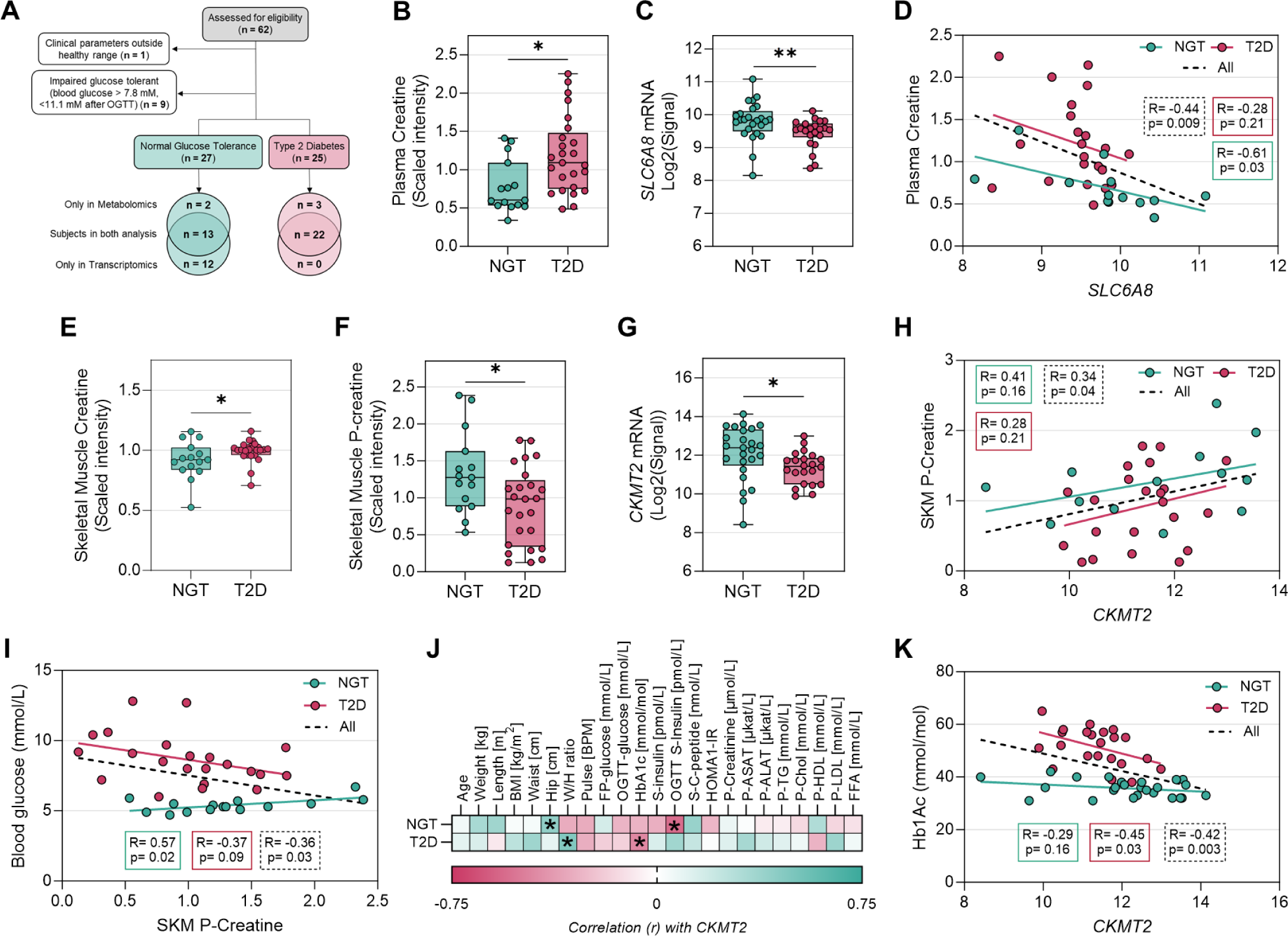
Skeletal muscle creatine metabolism is perturbed in men with type 2 diabetes and is linked with poorer glucose control. (A) Flow chart illustrating participant enrolment and analysis. Transcriptomic and metabolomic analysis shared samples from 13 individuals with NGT and 22 with T2D. (B) Plasma creatine levels from metabolomic analysis. (C) mRNA levels of the creatine transporter gene *SLC6A8* from microarray. (D) Pearson’s correlation between *SLC6A8* and plasma creatine levels. (E–F) Skeletal muscle (E) creatine and (F) phosphocreatine levels from metabolomic analysis. (G) mRNA levels of the sarcomeric mitochondrial creatine kinase gene *CKMT2* from microarray. (H) Pearson’s correlation between *CKMT2* and skeletal muscle phosphocreatine levels. (I) Pearson’s correlation between skeletal muscle phosphocreatine levels and fasting blood glucose. (J) Heatmap showing correlation between clinical characteristics and *CKMT2* expression. (K) Pearson’s correlation between *CKMT2* and Hb1Ac. (B) – (C) and (E) were analysed by Mann-Whitney test; (F) was analysed by Student’s t test; (G) was analysed by Welch’s t test. n= 15 NGT and 25 T2D in (B), (E) and (F); n= 25 NGT and 22 T2D in (C) and (G); n= 13 NGT and 22 T2D in (D), (H) and (K); n= 15 NGT and 22 T2D in (I). *, p<0.05; **, p<0.01. NGT, normal glucose tolerance; T2D, type 2 diabetes.

**Table 1.** Clinical characteristics of the subjects included in the metabolomic analysis.

|  | **Normal Glucose Tolerance** | **Type 2 Diabetes** | **P-value** |
| --- | --- | --- | --- |
| **Age (years)** | 60 ± 2.4 | 63 ± 1.3 | 0.1339 |
| **Weight (kg)** | 82 ± 2.4 | 88 ± 1.8 | 0.0306 |
| **Height (m)** | 1.8 ± 0.02 | 1.79 ± 0.01 | 0.4615 |
| **BMI (kg/m<sup>2</sup>)** | 25.95 ± 1.6 | 27.66 ± 0.47 | 0.0429 |
| **Waist (cm)** | 90 ± 1.6 | 100 ± 1.6 | 0.0003 |
| **Hip (cm)** | 97 ± 1.2 | 101 ± 1.2 | 0.0344 |
| **WHR ratio** | 0.93 ± 0.01 | 0.99 ± 0.01 | 0.0009 |
| **Pulse (BPM)** | 62 ± 2.0 | 68 ± 1.6 | 0.0242 |
| **FP-glucose (mmol/L)** | 5.41 ± 0.13 | 8.7 ± 0.37 | <0.0001 |
| **120-min-glucose (mmol/L)** | 6.05 ± 0.25 | 16.78 ± 0.58 | <0.0001 |
| **HbA1c (mmol/mol)** | 36.13 ± 1.13 | 51.19 ± 1.32 | <0.0001 |
| **HbA1c (%)** | 5.5 ± 0.10 | 6.8 ± 0.12 | <0.0001 |
| **S-insulin (pmol/L)** | 52.37 ± 5.30 | 83.6 ± 9.20 | 0.0182 |
| **120 min S-Insulin (pmol/L)** | 294.2 ± 43.41 | 327.5 ± 42.06 | 0.6155 |
| **P-Creatinine (μmol/L)** | 84.53 ± 2.65 | 80.72 ± 2.17 | 0.1809 |
| **P-ASAT (μkat/L)** | 0.42 ± 0.04 | 0.39 ± 0.03 | 0.6016 |
| **P-ALAT (μkat/L)** | 0.38 ± 0.06 | 0.48 ± 0.04 | 0.1318 |
| **P-TG (mmol/L)** | 1.08 ± 0.14 | 1.41 ± 0.13 | 0.0917 |
| **P-Chol (mmol/L)** | 5.27 ± 0.17 | 4.50 ± 0.14 | 0.0014 |
| **P-HDL (mmol/L)** | 1.33 ± 0.06 | 1.24 ± 0.06 | 0.3526 |
| **P-LDL (mmol/L)** | 3.47 ± 0.13 | 2.62 ± 0.14 | 0.0004 |
| **S-C-peptide (nmol/L)** | 0.69 ± 0.04 | 0.99 ± 0.08 | 0.0115 |
| **HOMA1-IR** | 1.71 ± 0.22 | 4.82 ± 0.65 | 0.0007 |

Results are mean ± SEM for individuals with Normal Glucose Tolerance (n=15) and Type 2 Diabetes (n=25) men. Statistical analysis was performed using Student's *t*-test.

**Table 2.** Clinical characteristics of the subjects included in the transcriptomic analysis.

|  | **Normal Glucose Tolerance** | **Type 2 Diabetes** | **P-value** |
| --- | --- | --- | --- |
| **Age (years)** | 58 ± 1.8 | 62 ± 1.3 | 0.0181 |
| **Weight (kg)** | 84 ± 1.7 | 89 ± 1.9 | 0.0577 |
| **Height (m)** | 1.80 ± 0.02 | 1.79 ± 0.01 | 0.6726 |
| **BMI (kg/m<sup>2</sup>)** | 25.98 ± 0.35 | 27.94 ± 0.48 | 0.0044 |
| **Waist (cm)** | 93 ± 1.4 | 101 ± 1.7 | 0.0030 |
| **Hip (cm)** | 99 ± 0.9 | 101 ± 1.3 | 0.2420 |
| **WHR ratio** | 0.94 ± 0.01 | 0.99 ± 0.01 | 0.0010 |
| **Pulse (BPM)** | 61 ± 1.4 | 68 ± 1.6 | 0.0064 |
| **FP-glucose (mmol/L)** | 5.46 ± 0.08 | 8.56 ± 0.39 | <0.0001 |
| **120-min-glucose (mmol/L)** | 5.89 ± 0.27 | 16.20 ± 0.69 | <0.0001 |
| **HbA1c (mmol/mol)** | 35.76 ± 0.64 | 50.44 ± 1.50 | <0.0001 |
| **HbA1c (%)** | 5.4 ± 0.06 | 6.8 ± 0.14 | <0.0001 |
| **S-insulin (pmol/L)** | 52.71 ± 5.58 | 92.58 ± 9.01 | <0.0001 |
| **120-min S-Insulin (pmol/L)** | 314.9 ± 50.8 | 350.5 ± 42.4 | 0.6439 |
| **P-Creatinine (µmol/L)** | 83.64 ± 2.36 | 79.71 ± 2.29 | 0.1426 |
| **P-ASAT (µkat/L)** | 0.38 ± 0.03 | 0.40 ± 0.03 | 0.7228 |
| **P-ALAT (µkat/L)** | 0.42 ± 0.05 | 0.50 ± 0.04 | 0.3181 |
| **P-TG (mmol/L)** | 1.25 ± 0.15 | 1.43 ± 0.13 | 0.2960 |
| **P-Chol (mmol/L)** | 5.40 ± 0.20 | 4.48 ± 0.15 | 0.0018 |
| **P-HDL (mmol/L)** | 1.30 ± 0.05 | 1.19 ± 0.06 | 0.1554 |
| **P-LDL (mmol/L)** | 3.56 ± 0.17 | 2.63 ± 0.15 | 0.0008 |
| **S-C-peptide (nmol/L)** | 0.69 ± 0.04 | 1.03 ± 0.08 | 0.0009 |
| **HOMA1-IR** | 1.86 ± 0.21 | 5.23 ± 0.65 | <0.0001 |

Results are shown as mean ± SEM for individuals with Normal Glucose Tolerance (n=25) and
Type 2 Diabetes (n=22) men. Statistical analysis was performed using Student's *t*-test.

### Glucose homeostasis is unaffected by changes in whole-body creatine levels

To investigate if changes in circulating and skeletal muscle creatine/phosphocreatine levels reflect or impair glucose homeostasis, male C57BL/6 mice were fed a high-fat diet for 8 weeks. Thereafter mice were injected with PBS, creatine, or the creatine analogue beta-guanidinopropionic acid (β-GPA, which acts as a competitive agonist to creatine for binding to the creatine transporter), for the last two weeks of the diet (**Figure 2A**). As expected, high-fat diet-fed mice had increased body weight and fat mass and reduced lean body mass compared with chow-fed mice, with no differences between treatments (**Figure 2B-D, Figure S2A-E**). Similarly, fasting glucose and glucose tolerance was unaffected by either creatine or β-GPA administration (**Figure 2E-G**). Consistently, both basal and insulin-stimulated glucose transport in soleus muscle was decreased in high fat diet-fed mice. Mice treated with β-GPA had improved basal glucose transport in soleus muscle and in insulin-stimulated glucose transport in EDL muscle (**Figure 2H-I**). Diet-induced insulin resistance and glucose intolerance was associated with skeletal muscle reduction in *Ckmt2* mRNA, thus recapitulating the results noted in people with type 2 diabetes (**Figure 2J**). Furthermore, despite the lack of changes in glucose metabolism, creatine administration altered the expression of key genes involved in creatine metabolism, oxidative stress, and substrate metabolism (**Figure 2K**). Notably, creatine treatment partially reverted the high-fat diet-induced downregulation of *Ckmt2* mRNA and upregulated the expression of *Hk2*, *Pdk4* and *Vegf* as well as genes encoding antioxidant enzymes (*Cat*, *Nos1*, *Nos3* and *Sod1*). These results provide evidence to suggest that the altered creatine metabolism observed in skeletal muscle in men with type 2 diabetes is not a cause of insulin resistance and type 2 diabetes.

**Figure 2.**
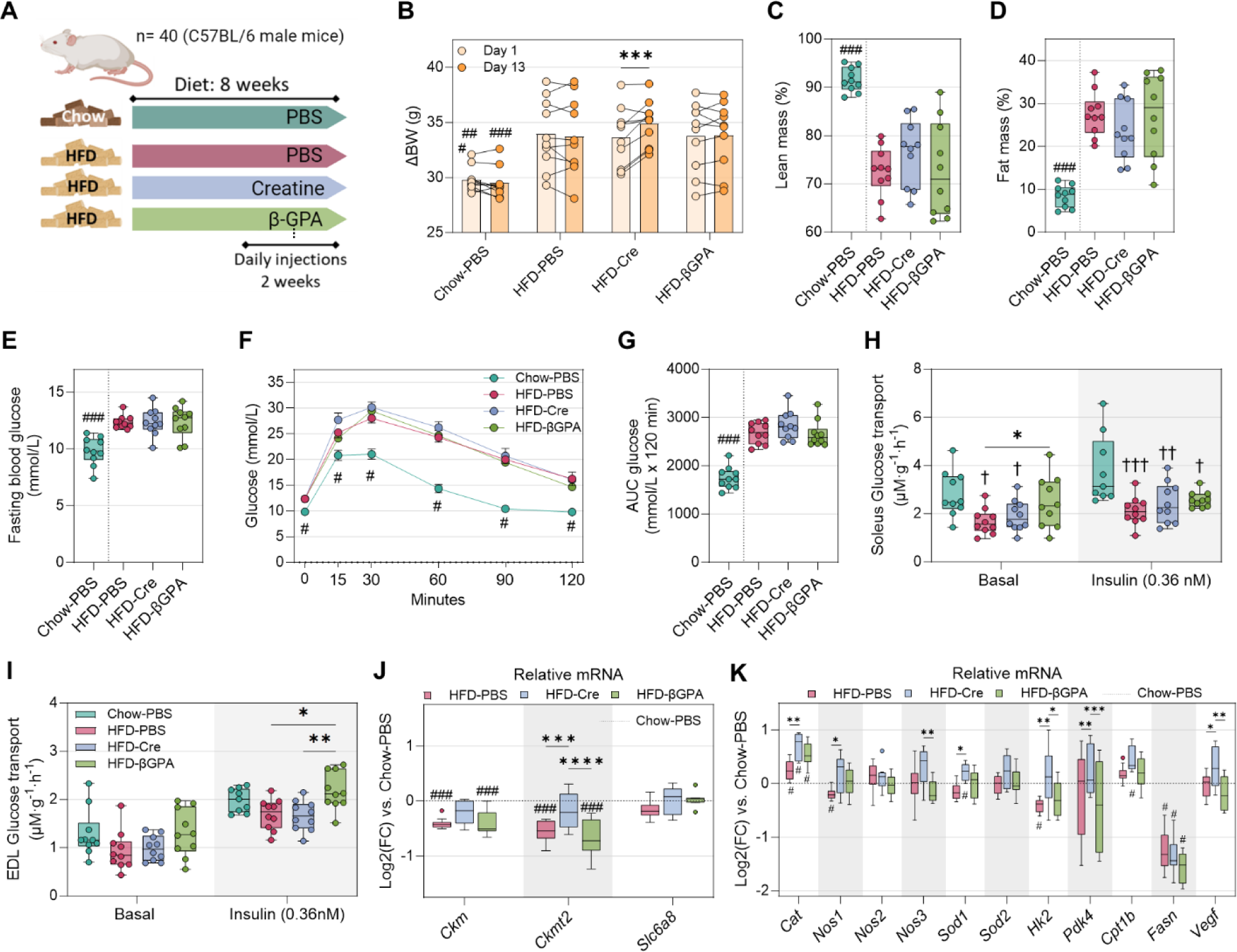
Alterations in creatine availability does not modulate glucose metabolism in high fat diet-fed mice. (A) Experimental design. (B) Body weight before starting intraperitoneal injections and at treatment day 13. (C – D) Percentage of (C) lean and (D) fat mass assessed by MRI at the end of the treatment. (E) Fasting blood glucose after treatment. (F) Time-course of an intraperitoneal glucose tolerance test performed at the end of treatment. (G) Area Under the Curve of the intraperitoneal glucose tolerance test. (H – I) (H) Ex-vivo soleus muscle and (I) EDL muscle glucose transport in basal and insulin-stimulated conditions. (J – K) Relative mRNA levels of genes involved in (J) creatine metabolism and in (K) in antioxidant defences and metabolism in the *tibialis anterior* muscle. (B) and (F) – (I) were analysed by two independent two-way ANOVA (mixed-effects model) followed by Dunnet’s post-hoc test (HFD groups vs. Chow-PBS) or Tukey’s post-hoc test (when comparing the different treatments within the HFD-fed mice). (C) – (E), (G) and (J-K) were analysed by two independent one-way ANOVA followed by Dunnet’s post-hoc test (HFD groups vs. Chow-PBS) or Tukey’s post-hoc test (when comparing the different treatments within the HFD-fed mice). (B) – (K), n=10 per group. *, p<0.05; **, p<0.01; ***, p<0.001; ****, p<0.0001; ###, p<0.001 vs. all HFD groups; †, p<0.05; ††, p<0.01; †††, p<0.001. Cre, Creatine; EDL, extensor digitorum longus; HFD, high fat diet.

### Silencing of *Ckmt2* impairs mitochondrial homeostasis in C2C12 myotubes

Next, we assessed the pathophysiological consequences of experimental reduction of skeletal muscle *Ckmt2* expression. We silenced *Ckmt2* in C2C12 myotubes with small interfering RNA, achieving a 60-90% knock-down at the mRNA level (**Figure 3A**) and a reduction in protein abundance (**Figure 3B**). Mitochondrial respiration was assessed in intact C2C12 myotubes by high resolution respirometry assays in an Oxygraph-2k (**Figure 3C**). *Ckmt2* silencing led to an overall decrease in mitochondrial respiration capacity, as evidenced by lower O_2_ flux during the ROUTINE (basal conditions with glucose available in media) and, notably, during the non-coupled state induced by FCCP, to reveal the capacity of the electron transfer system (ETS) (**Figure 3D**). The L/E ratio was increased in *Ckmt2*-silenced myotubes, most likely due to a lower ETS capacity (**Figure 3E**). To assess whether impaired mitochondrial respiration was associated with changes in ROS production, we used MitoSOX™ and Amplex™ fluorogenic dyes to measure superoxide (**Figure 3F**) and hydrogen peroxide (**Figure 3G**) production respectively. We found that *Ckmt2* silencing increased hydrogen peroxide in myotubes. Similarly, *Ckmt2* silencing led to a modest decrease in the mitochondrial membrane potential (**Figure 3H**), as assessed by quantifying the mitochondrial potential dye TMRE. Next, we incubated myotubes with D-[U-^14^C]-glucose and ^14^C-labelled CO_2_ measured to assess glucose oxidation via the TCA cycle. Consistent with the decrease in mitochondrial respiration, we found that *Ckmt2* silencing reduced glucose oxidation (**Figure 3I**). Interestingly, citrate synthase enzymatic activity – a well-known marker of mitochondrial content – was unaltered (**Figure 3J**), suggesting that the attenuation in mitochondrial function and oxidative phosphorylation was caused by intrinsic functional and/or structural changes rather than a reduction of mitochondrial numbers and density. Indeed, after analysing electron microscopy micrographs (**Figure 3K**) we found that *Ckmt2* silencing resulted in smaller mitochondria, without altering the shape or number (**Figure 3L-Q**).

**Figure 3.**
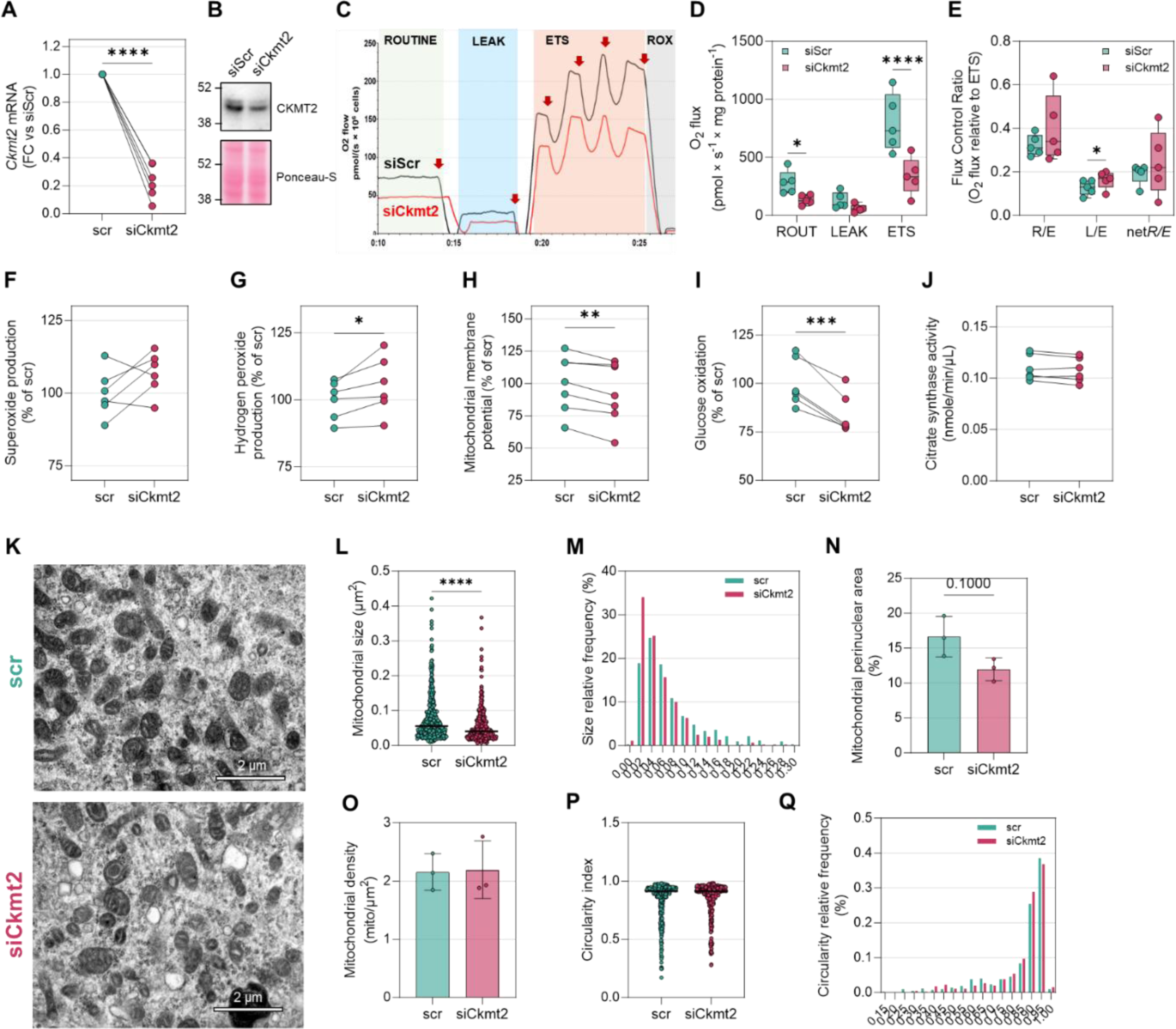
Silencing of *Ckmt2* impairs mitochondrial homeostasis in myotubes. (A) *Ckmt2* silencing efficiency at mRNA level in C2C12 myotubes. (B) Representative immunoblot of Ckmt2 silencing. (C) Representative high resolution respirometry assay conducted in intact C2C12 myotubes. Red arrows indicate sequential addition of compounds: oligomycin to assess leak respiration, FCCP titration to elicit maximal ETS respiration, and antimycin a to measure passive ROX oxygen consumption. (D) O2 flux during the three respiratory states. (E) O2 flux relative to the maximal respiration capacity. (F – H) Changes in (F) superoxide production, (G) hydrogen peroxide production, and (H) mitochondrial membrane potential assessed by fluorimetry. (I) Change in ^14^C radiolabelled glucose oxidation rate. (J) Citrate synthase activity assessed by enzymatic colorimetric assay. (K) Representative TEM images of C2C12 myotubes. (L – Q) Mitochondrial morphology parameters: (L) mitochondrial size and its (M) relative frequency distribution; (N) percentage of non-nuclear analysed area occupied by mitochondria; (O) mitochondrial density; and (P) mitochondrial circularity and (Q) its relative frequency distribution. (A) and (F) – (J) were analysed by a paired Student’s t test; (D) – (E) were analysed by two-way ANOVA (repeated measurements model) followed by Fisher’s LSD test; (L) and (N) – (P) were analysed by Mann-Whitney test. (A) – (J), n=5-7 per assay; (L) – (M) and (P) – (Q), n= 400 – 500 analysed mitochondria; (N) – (O), n=3.

### *Ckmt2* overexpression protects against lipid-induced metabolic stress in C2C12 myotubes independently of fatty acid handling

We investigated if *Ckmt2* overexpression protected against metabolic stress. C2C12 myotubes were transfected with a pCMV-Ckmt2 plasmid and, four days later, cells were incubated for 24 hours with either vehicle or 0.5 mM of oleate/palmitate to induced insulin resistance (**Figure 4A**). The pCMV-Ckmt2 vector effectively increased *Ckmt2* mRNA and protein (**Figure 4B-C**). Moreover, this expression remained elevated in oleate/palmitate-treated cells, while under control (empty vector) conditions palmitate treatment downregulated the expression of *Ckmt2* (**Figure 4D**). Overexpression of *Ckmt2* did not prevent the oleate/palmitate-induced downregulation of *Ppargc1a* (**Figure 4E**), suggesting that subsequent changes were not due to changes in the expression of this master regulator of mitochondrial function. While we did not detect lipid-induced alterations in *Sod1* mRNA levels (**Figure 4F**), *Ckmt2* overexpression prevented the decrease in *Sod2* mRNA levels induced by oleate/palmitate incubation (**Figure 4G**) and upregulated the expression of *Cat* (**Figure 4H**). Consistently, pCMV-Ckmt2 transfected myotubes were resistant to oleate/palmitate-induced activation of the stress-responsive p38 MAPK, as reflected by lower relative levels of T180/Y182 phosphorylation (**Figure 4I**). Notably, this apparent protection against lipid-induced stress was not associated with changes in oxidation of either palmitate or glucose (**Figure 4J-N**, **Figure S3A-B**), suggesting that the benefits derived from *Ckmt2* overexpression may be related to structural, rather than metabolic changes. Moreover, *Ckmt2* expression was particularly sensitive to oleate/palmitate incubation, resulting in a 55%reduction in mRNA levels as compared to myotubes exposed to BSA (**Figure S3C**).

**Figure 4.**
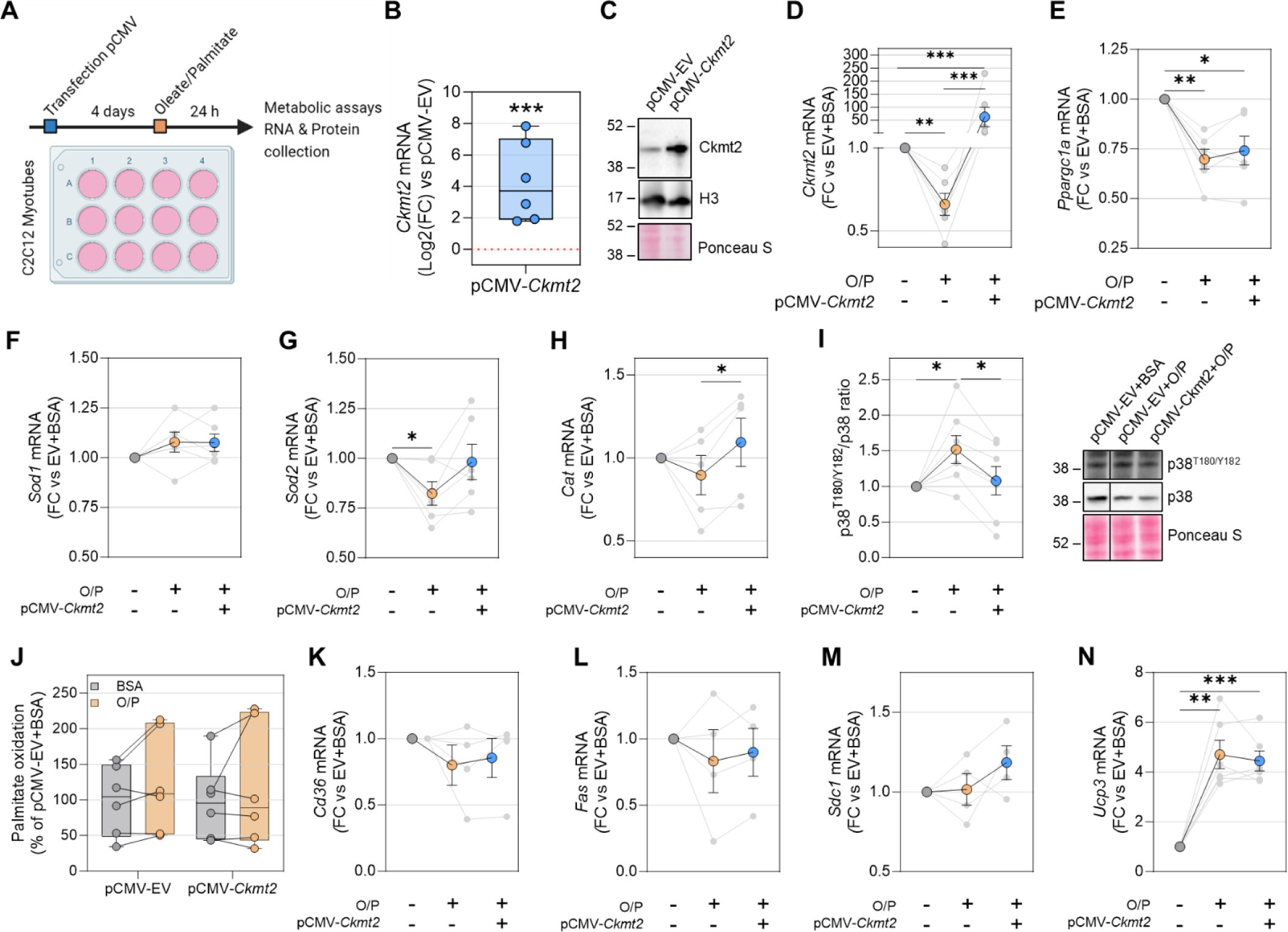
Overexpression of *Ckmt2* protects against lipid overload-induced metabolic stress. (A) Schematic experimental design. (B) Efficiency of pCMV-mediated *Ckmt2* overexpression in C2C12 myotubes. (C) Representative immunoblot of Ckmt2. (D – H) mRNA levels of (D) *Ckmt2*, (E) *Ppargc1a*, (F) *Sod1*, (G) *Sod2*, and (H) *Cat* on C2C12 myotubes incubated with oleate/palmitate and overexpression *Ckmt2*. (I) Ratio between activated p38^T180/Y182^ MAPK and total p38 MAPK content in C2C12 myotubes. A representative immunoblot is shown. (J) Palmitate oxidation capacity in myotubes overexpressing Ckmt2 and incubated with oleate/palmitate. (K – N) mRNA levels of genes involved in fatty acid handling, (K) *Sdc1*, (L) *Fas*, (M) *Sdc1*, and (N) *Ucp3*. (A) was analysed by a paired Student’s t test; (D) – (I) were analysed by one-way ANOVA followed by Tukey’s post-hoc test; (J) was analysed by two-way ANOVA. n=6 per condition except (K) – (M), where n=4. EV, empty vector; FC, Fold-change; O/P, oleate/palmitate.

### *Ckmt2* overexpression in skeletal muscle increases mitochondrial respiration and attenuates p38 MAPK activation in high fat diet-fed mice

To assess the potential protective role of *Ckmt2* overexpression against lipid-overload induced metabolic stress, C57BL/6 male mice were fed a high-fat diet for 11 weeks. Thereafter, the right and left *tibialis anterior* muscles were injected with pCMV-Ckmt2 or an empty vector, respectively, followed by metabolic analysis one week later (**Figure 5A-B**). Mice were injected with ^14^C-glucose and subjected to an oral glucose tolerance test and subsequently skeletal muscle was removed. Overexpression of *Ckmt2* did not alter skeletal muscle glucose uptake (**Figure 5C**) or phosphorylation of canonical proteins in the insulin-dependent signalling pathway (**Figure 5D**). In separate experiment, permeabilized fibre bundles from electroporated *tibialis anterior* muscles were used to perform high resolution respirometry assays (**Figure 5E**). *Ckmt2* overexpression increased Complex I-driven mitochondrial respiration and Complex I+II maximal respiration (**Figure 5F**), without altering the flux control ratio (**Figure 5G**). Protein abundance of the electron transport chain subunits was not altered in skeletal muscles electroporated with the pCMV-Ckmt2 vector (**Figure 5H**), suggesting ultrastructural remodelling. Moreover, *tibialis anterior* muscles electroporated with the pCMV-Ckmt2 plasmid had reduced p38 MAPK and Erk1/2 MAPK phosphorylation compared to the control leg (**Figure 5I**). These results indicate that *Ckmt2* overexpression may promote mitochondrial respiration and protect against the metabolic stress associated with lipid overload and insulin resistance.

**Figure 5.**
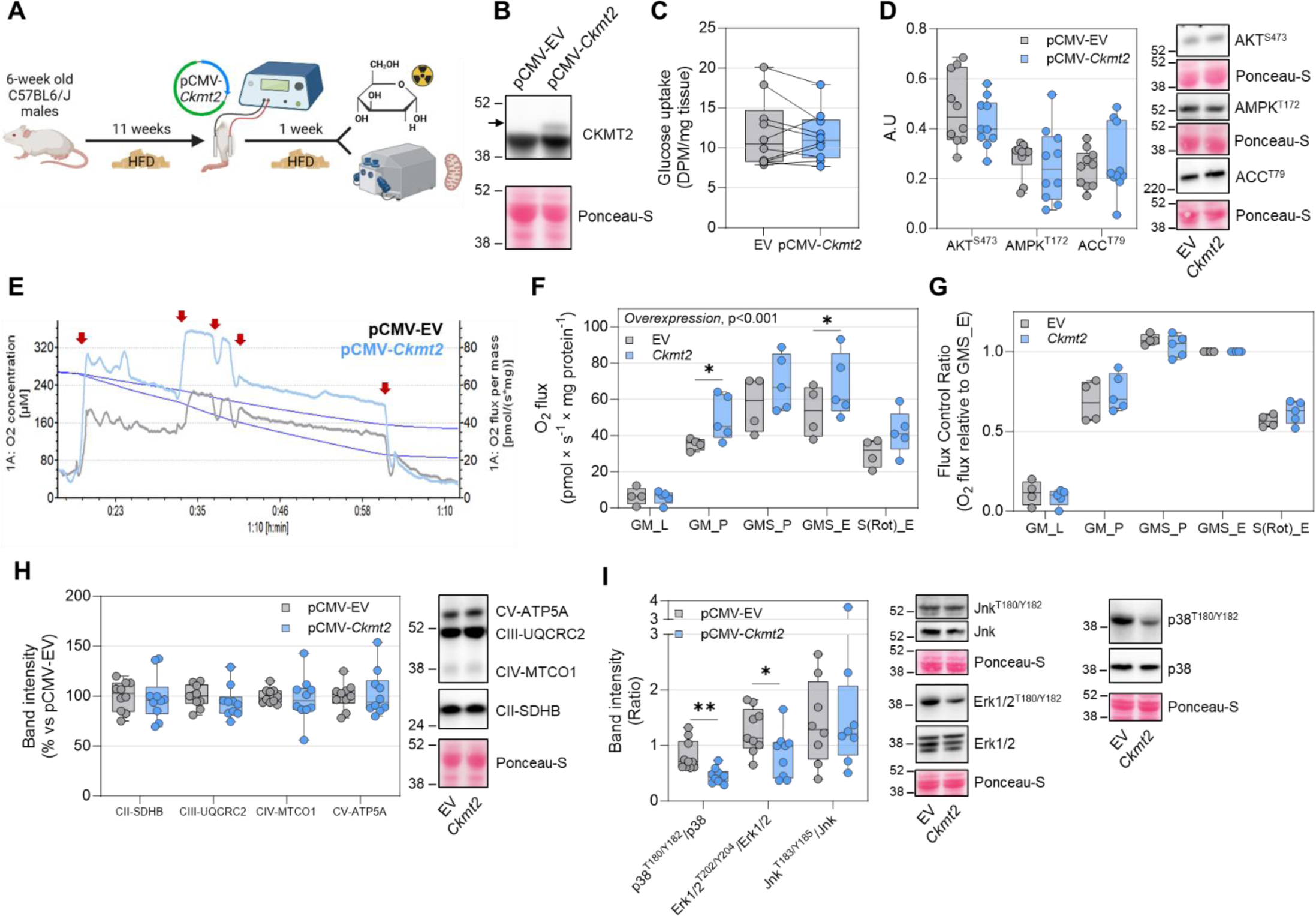
*In-vivo* overexpression of *Ckmt2* increases mitochondrial respiration and attenuates P38 activation in high fat diet-fed mice. (A) Scheme of the experimental design. (B) Representative immunoblot of Ckmt2 in *tibialis anterior* muscle homogenates. (C) Radiolabeled glucose uptake in tibialis anterior muscle. (D) Band densitometry and representative immunoblots of AKT^S473^, AMPK^T172^, and ACC^T79^ from tibialis anterior muscle. (E) Representative high resolution respirometry assay in permeabilized tibialis anterior muscle fiber bundles. Arrows indicate sequential addition of glutamate-malate-pyruvate (Complex I substrates), ADP, succinate (Complex II substrate), FCCP, rotenone and antimycin A. (F) Mitochondrial respirometry under different respiratory states. (G) Mitochondrial respiration normalized to maximal electron transfer system capacity. (H) Band densitometry and representative immunoblots of OXPHOS subunits. (I) Band densitometry and representative immunoblots of p38^T180/Y182^, Erk1/2^T202/Y204^, and Jnk^T183/Y182^ relative to their corresponding total protein levels. (C) – (D) and (H) – (I) were analyzed by paired Student’s t test, (F) and (G) were analysed by two-way ANOVA (mixed model) followed by Fisher’s LSD test. (C) – (D) and (H) – (I), n=10 per condition (except JNK in (I) where n=8); (F) and (G) n=4-5. EV, empty vector.

## DISCUSSION

Here we provide evidence linking type 2 diabetes to specific alterations in skeletal muscle creatine metabolism. Our findings indicate that elevated plasma creatine and reduced skeletal muscle phosphocreatine are markers, rather than drivers of peripheral insulin resistance. Furthermore, we elucidate the role of skeletal muscle CKMT2 in mitochondrial homeostasis in type 2 diabetes.

Increased plasma levels of creatine are linked to risk of developing type 2 diabetes^5,12^. Concordantly, we observed increased levels of circulating creatine in plasma samples from men with type 2 diabetes, as well reduced intramuscular phosphocreatine, recapitulating *in vivo* findings using phosphorus-31 magnetic resonance spectroscopy^13^. Plasma creatine levels were inversely correlated with expression of the creatine transporter (*SLC6A8*) in skeletal muscle, reinforcing the major contribution of this tissue to the systemic circulating levels of creatine. Conversely, in men with type 2 diabetes skeletal muscle phosphocreatine content was reduced and associated with impaired glucose homeostasis. Nevertheless, the effects of creatine loading on glucose homeostasis are inconclusive, with some reports showing glucose lowering effects^14,15^, while others report increased skeletal muscle glycogen storage with no systemic effects on glucose homeostasis^16,17^. Adding to this contention, creatine supplementation exacerbated glucose intolerance in a dexamethasone-induced muscle wasting rat model^18^. These findings raise the question as to whether creatine metabolism alterations are a cause or a consequence of insulin resistance and impaired glucose tolerance. Here we show that neither creatine supplementation, nor β-GPA-induced depletion of phosphocreatine modulate whole-body glucose metabolism in insulin resistant mice. Notably, β-GPA is an antihyperglycemic compound, activating AMPK signalling and GLUT4 translocation^19,20^. Consistently, β-GPA treatment increased glucose uptake in oxidative and glycolytic skeletal muscle. However, β-GPA-treated mice did not improve glucose tolerance or restored glycemia in insulin resistant mice. Improvements in specific metabolic parameters in β-GPA treated mice have been also reported without changes in glucose tolerance^21^, suggesting that dose, treatment duration, and underlying metabolic health, are critical factors influencing the metabolic outcomes associated with β-GPA^19,21,22^. Creatine treatment increased mRNA levels of antioxidant genes in mouse skeletal muscle without functional alterations in glucose metabolism, suggesting that creatine supplementation is not deleterious on insulin action. Collectively, our data demonstrate that changes in creatine and phosphocreatine levels are a consequence rather than a cause of insulin resistance.

Creatine is phosphorylated into phosphocreatine by creatine kinases. In skeletal muscle mitochondria, this reaction is catalysed by CKMT2^6^. Concomitant with changes in creatine metabolites, we also noted lower expression of *CKMT2* in skeletal muscle from men with type 2 diabetes. Indeed, CKMT2 protein has been found to be negatively enriched specifically in the mitochondrial proteome of patients with type 2 diabetes^23^. Moreover, eight weeks of high-fat diet feeding, which rendered mice insulin resistant, also recapitulated the lower skeletal muscle CKMT2 expression. The reduced muscle phosphocreatine in patients with type 2 diabetes might be reflective of low CKMT2 expression. Reduced phosphocreatine recovery rate is a conventional surrogate marker of impaired mitochondrial function^13^, which might partially reflect CKMT2 content in skeletal muscle. Indeed, we show that silencing of *Ckmt2* in C2C12 myotubes reduced basal and maximal mitochondrial respiration, suggesting that CKMT2 levels provides a molecular link between phosphocreatine and mitochondrial function. The depressed mitochondrial respiration in *Ckmt2-*silenced myotubes was accompanied by a moderate decrease in membrane potential. A hallmark of mitochondrial dysfunction is the loss of mitochondrial membrane potential, due to reduced ATP generation^24^, apoptotic signalling^25^ and disrupted mitochondrial dynamics^26^. Indeed, low membrane potential leads to the fragmentation of the mitochondrial network^26,27^, consistent with increased presence of smaller mitochondria in *Ckmt2*-silenced C2C12 myotubes. Overall, the alterations in mitochondrial homeostasis after *Ckmt2* silencing ultimately reduced basal glucose oxidation. Thus, CKMT2 plays a critical role in maintaining mitochondrial function and integrity, thereby arising as a potential candidate for mitochondrial dysfunction in type 2 diabetes.

We found that lipid overload downregulated *Ckmt2* expression, suggesting that intramuscular ectopic lipid accumulation contributes to the reduced abundance of CKMT2 in type 2 diabetes patients. Creatine kinases – and, particularly, CKMT2 – are extremely susceptible to oxidative damage^28,29^, which is often associated with exacerbated skeletal muscle lipid storage and lipid intermediates^30^. Thus, the lipid overload-induced reduction of CKMT2 plays a causal role in mitochondrial dysfunction and oxidative stress. We tested the hypothesis whether overexpression of CKMT2 could therefore protect against lipid-induced metabolic stress. Indeed, *Ckmt2* overexpression in either C2C12 myotubes or intact muscle mitigated the lipid-induced activation of p38 MAPK, a stress- and ROS-responsive kinase, implicated in the development of type 2 diabetes^31^. Moreover, *Ckmt2* overexpression upregulated the mRNA levels of the antioxidant genes *Sod2* and *Cat* in palmitate-exposed C2C12 myotubes, while in the *tibialis anterior* muscle oxidative phosphorylation was increased. We propose that CKMT2 prevents oxidative stress by upregulating antioxidant genes and promoting efficient mitochondrial respiration, thereby exerting a protection against metabolic stress associated with lipid overload. Thus, the CKMT2 reduction in skeletal muscle from people with type 2 diabetes constitutes a potential molecular link between lipid overload-mediated impairments of mitochondrial function and metabolism.

Changes in CKMT2 had profound effects on mitochondrial homeostasis and response to lipid-induced metabolic stress. However, mitochondrial content was unaltered, suggesting that CKMT2 could intrinsically modulate mitochondrial function. Of note, the *ex vivo* respirometry assays were conducted in the absence of creatine and phosphocreatine – potent modulators of mitochondrial respiration^32^ –, indicating that the observed increase in mitochondrial respiration was independent of concomitant changes in CKMT2 kinase activity and creatine-ADP stoichiometry. This result is compatible with CKMT1 isoform depletion-induced mitochondrial depolarization, independently of phosphocreatine availability, in HeLa cells ^33^. Moreover, both CKMT1 and CKMT2 induce and stabilize contact sites between the inner and outer mitochondrial membrane by colocalizing with ANT and VDAC^34–36^. Hence, we hypothesize that reduction of CKMT2 leads to mitochondrial membrane destabilization, subsequently impairing mitochondrial function and homeostasis, contributing to disruptions in energy metabolism observed in metabolic disorders (**Figure 6**).

**Figure 6.**
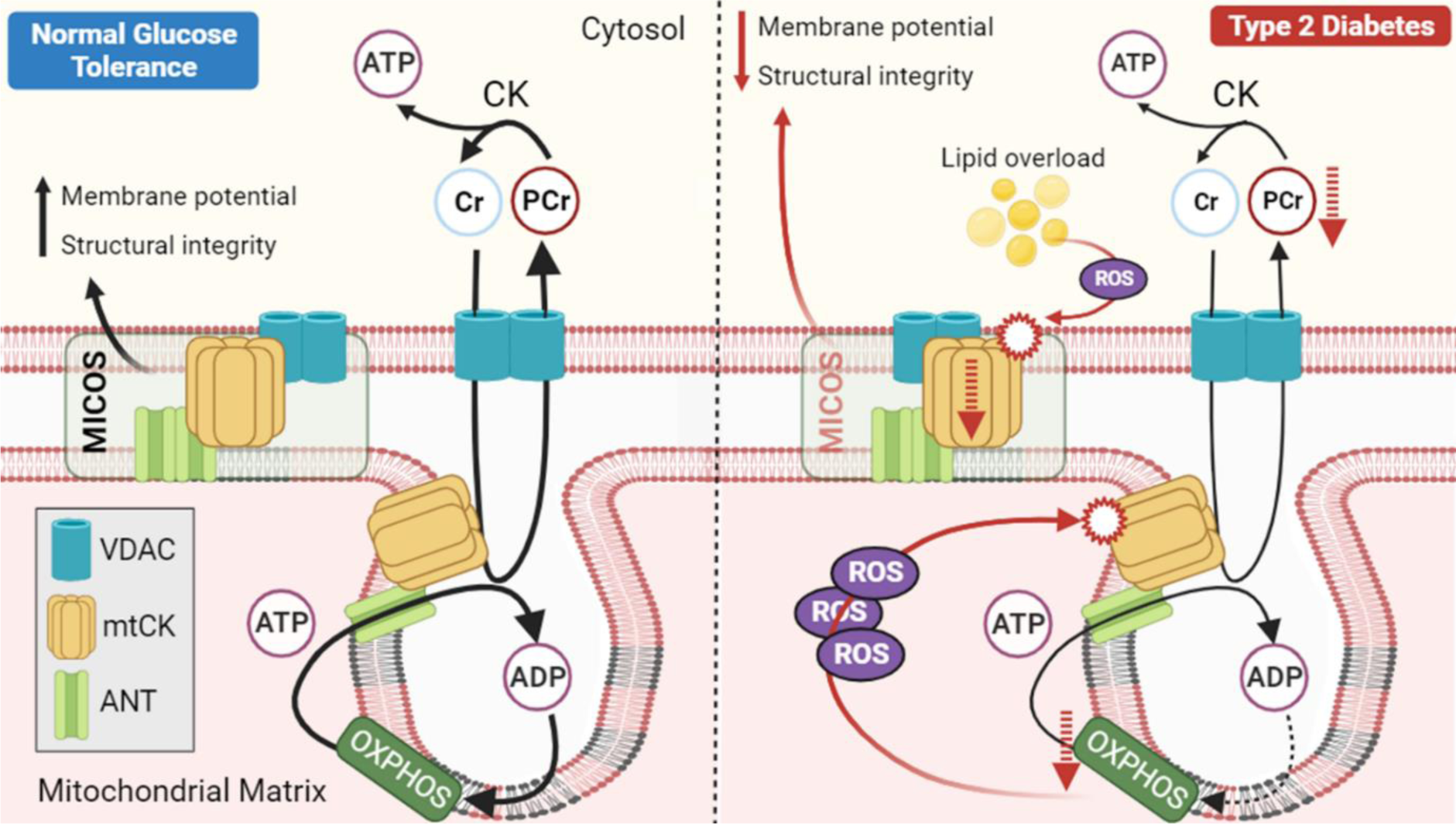
Proposed mechanism for the implication of reduced CKMT2 expression in mitochondrial function in type 2 diabetes skeletal muscle. CKMT2 participates in the formation of mitochondrial contact sites (MICOS) that contribute to the structural integrity of the mitochondrial membrane and regulate the membrane potential. Moreover, by localising in close proximity of the electron transport chain, CKMT2 hydrolyses ATP derived from oxidative phosphorylation (OXPHOS) to convert creatine into phosphocreatine and releasing ADP, thus being an important modulator of mitochondrial respiration. We propose that, in type 2 diabetes, intramuscular lipids and concomitant production of reactive oxygen species (ROS) damage and downregulate CKMT2. Lower CKMT2 content leads to decreased membrane potential, integrity and mitochondrial respiration while exacerbating ROS production and lowering glucose oxidation capacity.

Altered mitochondrial homeostasis in skeletal muscle is regarded as a hallmark of type 2 diabetes and insulin resistance^37,38^, although the direction of causality is a matter of intense debate^39–41^. The effects of either reduced or increased CKMT2 as reported here were evident from experiments performed in the absence of insulin, thereby disassociating mitochondrial homeostasis from insulin signalling. Thus, the reduced CKMT2 expression in skeletal muscle from patients with type 2 diabetes likely contributes to impaired mitochondrial function, providing a plausible mechanism that disassociates mitochondrial capacity from insulin sensitivity.

In conclusion, CKMT2 plays a key role in skeletal muscle mitochondrial homeostasis. Ckmt2 reduction recapitulates key features of the skeletal muscle dysfunction in type 2 diabetes, including impaired mitochondrial function, increased ROS production and reduced glucose metabolism. Conversely, Ckmt2 overexpression increases oxidative phosphorylation and protects against lipid-induced metabolic stress. Collectively, we reveal a previously underappreciated role for CKMT2 on mitochondrial homeostasis, independent of creatine metabolism and insulin action. Thus, therapeutic strategies to modulate CKMT2 expression may be efficacious in the treatment of metabolic disease.

## AKNOWLEDGMENTS

DRR was supported by a Novo Nordisk postdoctoral fellowship run in partnership with Karolinska Institutet. This study was supported by the EFSD/Lilly Young Investigator Research Award Programme. MR was supported by grants from Margareta af Uggla’s foundation, the Swedish Research Council, ERC-SyG SPHERES (856404), the NovoNordisk Foundation (including the MeRIAD consortium Grant number 0064142), Knut and Alice Wallenbergs Foundation (including Wallenberg Clinical Scholar), CIMED, the Swedish Diabetes Foundation, the Stockholm County Council, and the Strategic Research Program in Diabetes at Karolinska Institutet.

We acknowledge Ann-Marie Pettersson for her expertise and support in conduction animal experiments.

Figure schematics were created with BioRender.com.

## AUTHOR CONTRIBUTIONS

Conceptualization, D.R-R., M.R., J.R.Z., and A.K.; investigation, D.R-R., D.S.P.S.F.G., L.A.P., N.D.L., A.V.C., and S.M, formal analysis, D.R-R. and D.S.P.S.F.G.; data curation, D.R-R.; funding acquisition, D.R-R., M.R., J.R.Z. and A.K.; methodology, D.R-R., D.S.P.S.F.G. and A.V.C.; resources, E.N., J.R.Z. and A.K.; supervision, J.R.Z. and A.K.; Visualization, D.R.-R.; writing – original draft, D.R-R.; writing – review and editing, D.R-R., D.S.P.S.F.G., L.A.P., N.D.L., A.V.C., S.M., E.N., M.R., J.R.Z. and A.K.

## METHODS

### Participants

The study was approved by the regional ethical review board in Stockholm and procedures were conducted according to the Declaration of Helsinki. Informed written consent was obtained from each participant. After clinical health screening and application of exclusion criteria (Figure 1A), twenty-seven men with normal glucose tolerance and twenty-five men with type 2 diabetes were included in this study. Participants with type 2 diabetes were treated with the following glucose-lowering drugs: metformin (n = 20; daily dose range 500–3000 mg) and/or sulfonylurea (glibenclamide [glyburide], n = 5, 2–5 mg daily; glipizide, n = 1, 5 mg daily). Glucose lowering medication was taken by the individuals with type 2 diabetes after the skeletal muscle biopsy collection.

Participants arrived at the clinic after an overnight 12 h fast. Fasting blood samples were obtained. After subcutaneous application of local anaesthesia (lidocaine hydrochloride 5 mg/ml; Aspen Nordic, Denmark), vastus lateralis biopsies were obtained using a conchotome (AgnTho’s, Sweden). Biopsies were immediately snap-frozen in liquid nitrogen and stored at −80°C. Total RNA Microarray analysis was performed with the Affymetrix GeneChip Human Transcriptome Array 2.0 (Thermo Fisher Scientific, MA). Metabolomic analysis was performed by Metabolon (Durham, NC, USA) as described^42^.

Clinical characteristics are of the individuals included in the metabolomic and transcriptomic analyses are shown in **Table 1** and **Table 2**.

### Mouse studies

All mouse experiments were carried out in C57BL/6 male mice purchased from Charles River (Germany). Mice were housed in groups of 4 in ventilated cages with a 12 h light/12 h dark cycle in a temperature-controlled (20-24°C) environment with ad libitum access to food and water. Mice were purchased and housed as described above, with ad libitum access to food and water. All experimental procedures were approved by the Stockholm North Animal Ethical Committee (ethical permit N38/15) and carried out according to the animal care guidelines European Union laws.

#### Creatine analogues study

At 6 weeks of age, mice were fed either a standard chow diet (n=10) (4% kcal from fat, R34; Lantmännen) or a 60% fat diet (n=30) (Research Diet, New Brunswick, NJ) for 8 weeks. At 12 weeks of age, mice were randomized to the treatment groups (n = 10/group) based on body weight and received daily intraperitoneal injections of PBS (20 ml/kg) or creatine (0.5 mg/kg body weight) for 14 days; or GPA (0.4 mg/kg body weight) for 1 week (Figure 2A). At the end of the treatment, body composition was assessed by MRI (EchoMRI-100, EchoMRI™).

#### Intraperitoneal glucose tolerance test

One day before the last injection, mice were fasted for 4 hours and glucose tolerance was determined. Glucose (2 g/kg body weight) was intraperitoneally administered, and blood glucose levels were monitored for 2 hours with a glucometer (Contour XY, Bayer) using tail vein blood samples.

#### Electroporation-induced Ckmt2 overexpression experiments

At 6 weeks of age, mice were fed a 60% fat diet (n=15) (Research Diet, New Brunswick, NJ) for 11 weeks before the electroporation procedure. Briefly, mice were anesthetized with 2% isoflurane anaesthesia and both hindlegs were shaved and cleaned with chlorhexidine. Hyalounidase (30 µl of 1 unit/µl) was injected intramuscularly in the tibialis anterior muscle and mice returned to their homecage. After 2 hours, mice were anaesthetized again with 2% isoflurane and the tibialis anterior muscle of each leg was injected with either a pCMV plasmid encoding *Ckmt2* or an empty vector. Muscles were electroporated immediately after by delivering 220 V/cm in 8 pulses of 20 milliseconds using an ECM 830 electroporator (BTX Harvard Apparatus, Holliston, MA)^43^.

#### Modified glucose tolerance test and in vivo glucose uptake

The effects of *Ckmt2* overexpression on skeletal muscle glucose transport was assessed with a modified glucose tolerance test. Briefly, a standard 2-h oral glucose tolerance test was performed seven days after electroporation. Fifteen minutes after oral gavage of glucose (3 g/kg body weight), mice were intraperitoneally injected with radioactive ^3^H-2-Deoxy-D-glucose (ARC #ART0103; 1mCi/mL; SA: 60Ci/mmol). At the end of the test (120 min), mice were rapidly anaesthetised, and the tibialis anterior muscles were immediately dissected and frozen in liquid nitrogen.

Tibialis anterior skeletal muscles were homogenized in 300 µl of PI3K buffer (137 mM NaCl, 2.7 mM KCl, Tris pH 7.8, 1 mM MgCl_2_, 1% Triton X-100, 10% Glycerol, 10 mM NaF, 1 mM EDTA), rotated at 4°C for 1 hour, and centrifuged at 13000 xg for 10 min at 4°C. Thirty microliters of supernatants were analysed by liquid scintillation counting to determine accumulation of ^3^H-2-Deoxy-D-glucose, and the remaining portion was stored at −80 °C for immunoblot analysis.

### Cell studies

#### Culture of the murine C2C12 muscle cell line

C2C12 myoblasts were cultivated in Dulbecco’s Modified Eagle’s Medium (DMEM) supplemented with 10% calf serum and containing 25 mM glucose, 4 mM L-glutamine and 1 mM sodium pyruvate. Myogenic differentiation was induced in fully confluent cells by switching the calf serum in the growth media for 2% horse serum. The experiments were performed after 5 days of differentiation.

#### Silencing and overexpression of Ckmt2

Cells were transfected with 200 nM of siRNA targeting CKMT2 (siCkmt2) or scrambled siRNA (siScr) as a control following Lipofectamine RNAiMax manufacturer’s instructions. For overexpression, cells were transfected with a pCMV6-Entry vector containing the sequence of *Ckmt2* or an empty pCMV6-Entry vector as control following Lipofectamine 3000 manufacturer’s instructions. Transfection and differentiation media switch were performed simultaneously.

#### Treatment with palmitate/oleate

Fatty acid-free BSA was solubilized in differentiation media supplemented with 10 mM HEPES by agitation at 55°C for 15 minutes. Oleate and palmitate were added to a final concentration of 0.5 mM each and incubated at 55°C for 15 minutes. The solution was cooled down and filter sterilized (0.22 µm) before adding it to the cells. Control cells were treated with a vehicle consisting of 1% ethanol and 0.14% DMSO in HEPES-supplemented differentiation media. Myotubes were treated on the 5th day of differentiation for 18 hours.

### RNA extraction, cDNA synthesis and quantitative PCR

RNA was extracted with TRIzol according to the manufacturer’s instructions. RNA quality was assessed by spectrophotometry and 1 µg was used as input for cDNA synthesis using the High-Capacity cDNA Reverse Transcription Kit. cDNA was diluted and 25 ng were used in qPCR with the indicated primer pairs (**Supplemental Table 1**). Relative mRNA quantification was calculated and normalized by RPL39 (cell studies) or HPRT (mouse studies) gene expression.

### Western Blot

Skeletal muscle tissue was processed as described in the electroporation procedure. Cells were collected in RIPA buffer (Thermo Scientific) and homogenized with the aid of a sonicator. Insoluble material was sedimented by centrifugation at 16000 xg and protein concentration in the supernatant was determined by Bradford assay. Laemmli buffer was added to the samples and the proteins were resolved by gradient SDS-PAGE (4-12%). Wet transfer was used to blot the proteins into PVDF membranes, which were then stained with Ponceau S for normalization purposes and quality control. The membranes were blocked with 5% skimmed milk in TBS-T (20 mM Tris-HCl, 150 mM NaCl, 0.02% (v/v) Tween 20, pH 7.5) for 1 hour. The membranes were incubated overnight with primary antibody as indicated and, after several washes, incubated for 1 hour with HRP-conjugated secondary antibodies. Specific proteins were detected by chemiluminescence (ECL) in an imager (Gel Doc XR System, BioRad). Protein bands were quantified by densitometry using ImageLab software (BioRad) and normalized by the Ponceau S densitometric volume of the respective gel lane.

### ROS production assays

Cells were seeded in 96-well clear bottom black plates and treated as indicated. Mitochondrial superoxide content was measured by MitoSOX probe fluorescence following manufacturer instructions. Briefly, cells were incubated with 5 µM MitoSOX in growth media without serum for 30 minutes at 37°C and washed with PBS. Fluorescence at 510 nm/580 nm (ex./em.) was measured in a plate reader. Extracellular hydrogen peroxide content was measured with Amplex UltraRed Krebs-Henseleit Buffer in the presence of 0.2 mU/mL horseradish peroxidase at 37°C. After an initial 10 minutes of incubation, fluorescence at 540 nm/590 nm (ex./em.) was measured every 15 minutes for 90 minutes, and the rate of fluorescence increase after background removal was calculated.

### High resolution respirometry

#### C2C12 myotubes

C2C12 myotubes were differentiated in a 6-well plate. The day the experiment, cells were dissociated by adding 100-150 µl of trypsin and resuspended in 4 ml of warmed DMEM. 1-2 x 10^6^ cells were transferred to the respiration chambers. An aliquot of the cells was kept for later protein quantification and further normalization.

The effects of *Ckmt2* silencing on mitochondrial respiration was assessed by performing a Phosphorylation Control Protocol with Intact Cells^44^ in an Oxygraph 2k (Oroboros). Mitochondrial O_2_ consumption is measured in basal conditions (ROUTINE, cells suspended in DMEM), under oligomycin-induced (2µ-ml) ATP synthase inhibition (LEAK respiration) and under maximal respiration after FCCP titration in 1 µM steps (ETS, electron transfer system capacity). These measurements were corrected by the passive, non-mitochondrial O_2_ consumption obtained after the addition of the Complex III inhibitor Antimycin (2.5 μM) (ROX).

#### Skeletal muscle fibre bundles

Fiber bundles from electroporated tibialis skeletal muscle anterior were isolated and permeabilized with saponin (5 mg/ml) in ice-cold biopsy preservation solution BIOPS (10 mM Ca-EGTA buffer, 0.1 µM free calcium, 20 mM imidazole, 20 mM taurine, 50 mM K-MES, 0.5 mM DTT, 6.56 mM MgCl_2_, 5.77 mM ATP, 15 mM phosphocreatine) as described elsewhere^45^. Fiber bundles (1.5 – 2.4 mg) were added in the respiratory chambers containing 2.2 ml of mitochondrial respiration medium MIR05 (110 mM sucrose, 20 mM HEPES, 20 mM taurine, 60 mM K-lactobionate, 3 mM MgCl_2_, 10 mM KH_2_PO_4_, 0.5 mM EGTA, 1 g/l BSA, pH 7.1). After adding the fibres into the respiration chamber, the chamber was closed, oxygen concentration was increased to ∼400 µM and allowed it to equilibrate before adding any substrates. Complex I leak respiration was assessed by adding 2 mM malate, 10 mM pyruvate and 10 mM glutamate. After O_2_ consumption rates were stabilized, ADP was added to a final concentration of 5 mM, triggering Complex I-driven oxidative phosphorylation (OXPHOS). Complex II respiration was stimulated by adding 10 mM succinate, obtaining the OXPHOS capacity of Complex I + Complex II. ETS maximal capacity was determined by titrating carbonyl cyanide-4-(trifluoromethoxy) phenylhydrazone (FCCP) in 0.5 μM steps. Complex II-linked ETS capacity was measured by inhibiting Complex I with 0.5 µM rotenone. These measurements were corrected by ROX after adding 2.5 μM antimycin A. O_2_ consumption rates were normalized to the fibre bundle weight added in each assay, Mitochondrial membrane integrity and damage was assessed by adding cytochrome c after measuring Complex I respiration and observing no differences in O_2_ consumption.

### Citrate synthase activity assay

Cells were seeded in 96-well clear plates. Citrate synthase activity in C2C12 myotubes was measured with a Citrate Synthase Activity Assay Kit (MAK193, Sigma-Aldrich) and following the manufacturer’s instructions. Absorbance at 412 nm was read every 5 minutes for 40 minutes.

### Mitochondrial membrane potential

Cells were seeded in 96-well clear bottom black plates. Mitochondrial membrane potential in C2C12 myotubes was measured in a fluorescence plate reader (535 nm/580 nm (ex./em, respectively) with 50 nM TMRE and following the TMRE Mitochondrial Membrane Potential Assay Kit (ab113852, Abcam) manufacturer’s instructions.

