## Supplemental material for "Decreased sarcomeric mitochondrial creatine kinase 2 impairs skeletal muscle mitochondrial function independently of insulin action in type 2 diabetes"

### SUPPLEMENTAL INFORMATION

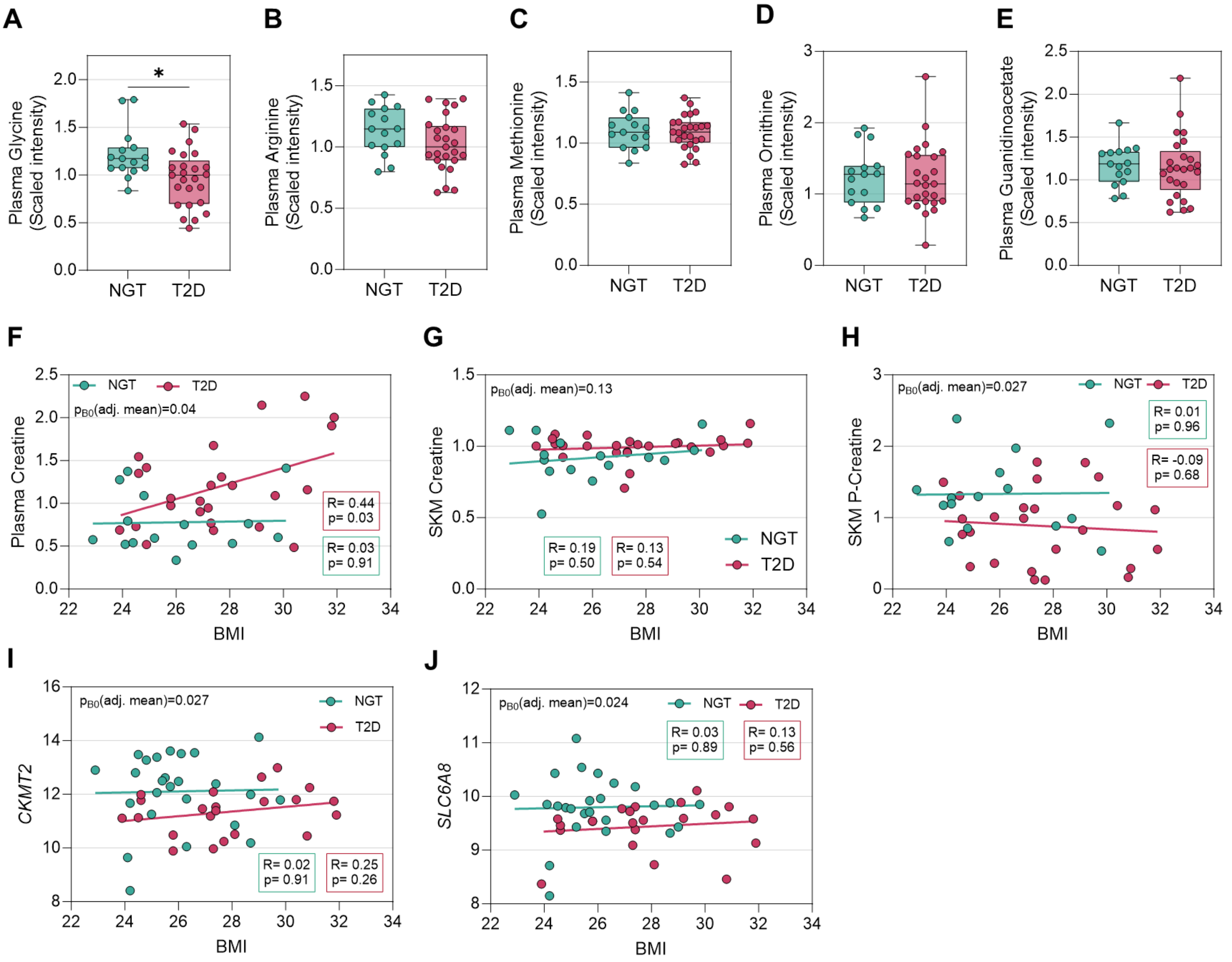


**Supplemental Figure 1 (supporting data to Figure 1)**

(A – E) Plasma levels of creatine precursors from metabolomic analysis.

(F – J) Simple linear regression analysis between BMI and creatine metabolites and genes displaying the intercept $B_0$ (ANCOVA adjusted mean) p-value.

(A) and (D) were analysed by Mann-Whitney test; (B), (C) and (E) were analysed by Student's t test. (A) – (H), n= 15 NGT and 25 T2D; (A) – (H), n= 25 NGT and 22 T2D. *, p<0.05. NGT, normal glucose tolerance; T2D, type 2 diabetes NGT, normal glucose tolerance; T2D, type 2 diabetes.

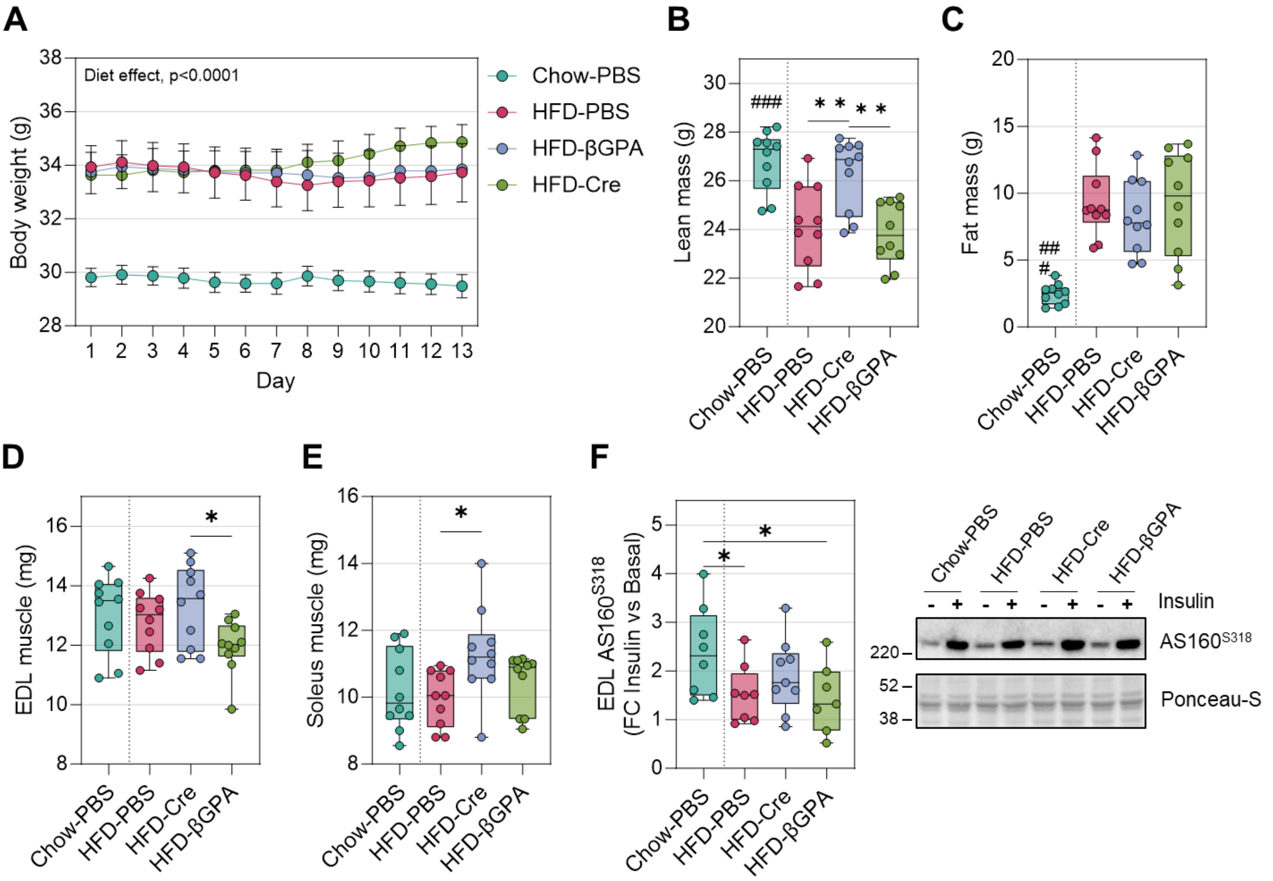


**Supplemental Figure 2 (supporting data to Figure 2)**

(A) Body weight across treatment period.

(B) Lean mass assessed by MRI at the end of the treatment.

(C) Fat mass assessed by MRI at the end of the treatment.

(D) EDL muscle weight after dissection.

(E) Soleus muscle weight after dissection.

(F) Fold-change of band densitometry and representative blot of $AS^{S318}$ from basal and insulin-stimulated EDL muscles.

(A) was analysed by two independent two-way ANOVA (mixed-effects model) followed by Dunnet's post-hoc test (when comparing each HFD group vs. Chow-PBS) or Tukey's post-hoc test (when comparing the different treatments within the HFD-fed mice). Significance for each individual timepoint not shown. (B) – (F) were analysed two independent one-way ANOVA followed by Dunnet's post-hoc test (when comparing each HFD group vs. Chow-PBS) or Tukey's post-hoc test (when comparing the different treatments within the HFD-fed mice). (A) – (E), n=10; (F), n=7-9. Cre, Creatine; EDL, extensor digitorum longus; HFD, high fat diet.

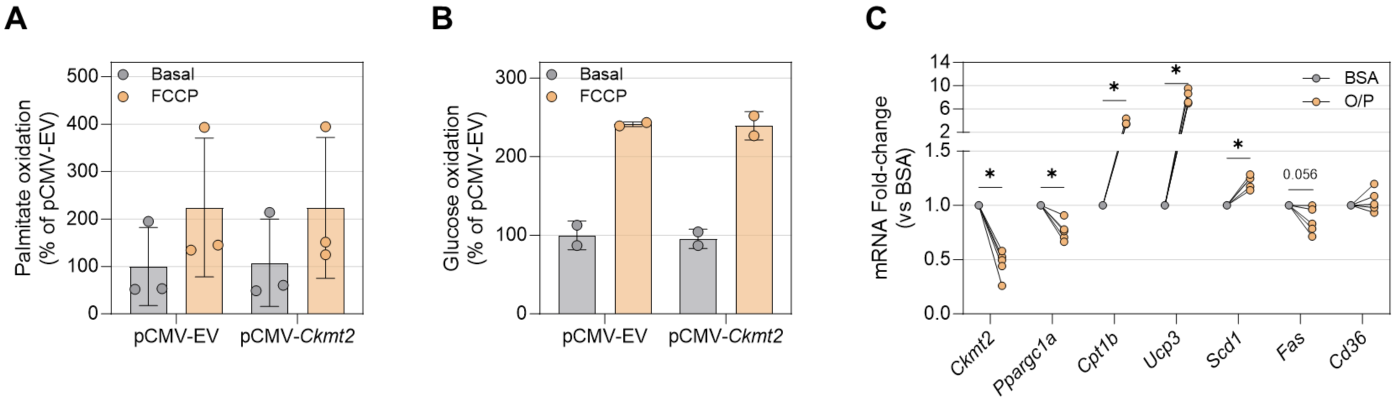


**Supplemental Figure 3 (supporting data to Figure 4)**

(A) Effects of FCCP on palmitate oxidation on C2C12 myotubes overexpressing *Ckmt2*.

(B) Effects of *Ckmt2* overexpression on glucose oxidation on C2C12 myotubes.

(C) Effects of incubation of oleate/palmitate in non-transfected C2C12 myotubes on the mRNA levels of *Ckmt2*, *Ppargc1a*, and genes involved in lipid metabolization.

(A) and (B) were analysed by two-way ANOVA. n=2-3; (C) was analysed by paired Student's t test (n=4). FC, Fold-change; O/P, oleate/palmitate.

**Table S1. Primer sequences used in the analysis of relative mRNA levels.**

| Gene | Forward sequence (5' to 3') | Reverse sequence (5' to 3') |
|---|---|---|
| **Cat** | AGCGACCAGATGAAGCAGTG | TCCGCTCTCTGTCAAAGTGTG |
| **Ckm** | CTGACCCCTGACCTCTACAAT | GAAGGGGTGACCTGGGTTG |
| **Ckmt2** | ACACCCAGTGGCTATACCCTG | CCGTAGGATGCTTCATCACCC |
| **Cpt1b** | CCCATGTGCTCCTACCAGAT | CGAGGATTCTCTGGAACTGC |
| **Fas** | CCCTTGATGAAGAGGGATCA | ACTCCACAGGTGGGAACAAG |
| **Hk2** | AGAGAACAAGGGCGAGGAG | GGAAGCGGACATCACAATC |
| **Hprt** | ACAGGCCAGACTTTGTTGGA | ACTTGCGCTCATCTTAGGCT |
| **Nos1** | GACTGATGGCAAGCATGACTTC | GCCCAAGGTAGAGCCATCTG |
| **Nos2** | TGACGGCAAACATGACTTCAG | GCCATCGGGCATCTGGTA |
| **Nos3** | TCTGCGGCGATGTCACTATG | CCATGCCGCCCTCTGTT |
| **Pdk4** | GGATTACTGACCGCCTCTTTAG | GTAACCAAAACCAGCCAAAGG |
| **Ppargc1a** | ACCAGTACAACAATGAGCCTGCGA | TCCAGTGTCTCTGTGAGAACCGC |
| **Rpl39** | CAAAATCGCCCTATTCCTCA | AGACCCAGCTTCGTTCTCCT |
| **Slc6a8** | TCCTGGCACTCATCAACAG | ATGAAGCCCTCCACACCTAC |
| **Scd1** | TGCGATACACTCTGGTGCTC | TAGTCGAAGGGGAAGGTGTG |
| **Sod1** | TGGTGGTCCATGAGAAACAA | GTTTACTGCGCAATCCCAA |
| **Sod2** | CAGACCTGCCTTACGACTATGG | CTCGGTGGCGTTGAGATTGTT |
| **Vegf** | CTGCTGTAACGATGAAGCCCTG | GCTGTAGGAAGCTCATCTCTCC |
| **Ucp3** | GACCCACGGCCTTCTACAAA | TCAAAACGGAGATTCCCGCA |
